# Optogenetic activation of basolateral amygdala projections to nucleus accumbens core promotes cue-induced expectation of reward but not instrumental pursuit of cues

**DOI:** 10.1101/2021.10.20.465037

**Authors:** Alice Servonnet, Pierre-Paul Rompré, Anne-Noël Samaha

## Abstract

Reward-associated conditioned stimuli (CS) can acquire predictive value, evoking conditioned approach behaviors that prepare animals to engage with forthcoming rewards. Such CS can also acquire conditioned reinforcing value, becoming attractive and pursued. Through their predictive and conditioned reinforcing properties, CS can promote adaptive (*e.g*., locating food) but also maladaptive responses (*e.g*., drug use). Basolateral amygdala neurons projecting to the nucleus accumbens core (BLA→NAc core neurons) mediate the response to appetitive CS, but the extent to which this involves effects on the predictive and/or conditioned reinforcing properties of CS is unclear. Thus, we examined the effects of optogenetic stimulation of BLA→NAc core neurons on conditioned approach behavior and on the instrumental pursuit of a CS, the latter a measure of conditioned reinforcement. Water-restricted, adult male rats learned that a light-tone compound cue (CS) predicts water delivery. Pairing optogenetic stimulation of BLA→NAc core neurons with CS presentation potentiated conditioned approach behavior, and did so even under extinction conditions, when water was omitted. This suggests that BLA→NAc core neurons promote cue-induced expectation of rewards. Rats also received instrumental conditioning sessions during which they could lever press for CS presentations, without water delivery. Optogenetic stimulation of BLA→NAc core neurons either during these instrumental test sessions or during prior CS-water conditioning did not influence lever responding for the CS. This suggests that BLA→NAc core neurons do not influence the conditioned reinforcing effects of CS. We conclude that BLA→NAc core neurons promote cue-induced control over behavior by increasing cue-triggered anticipation of rewards, without influencing cue ‘wanting’.

**SIGNIFICANCE STATEMENT:** Environmental cues associated with rewards guide animals toward rewards essential for survival such as food. Reward-associated cues can also evoke maladaptive responses such as in addiction. Reward cues guide behavior in two major ways. First, they evoke approach toward imminent rewards, preparing animals to engage with these rewards. Second, cues can become attractive themselves, such that animals will learn new behaviors simply to obtain them. Here we show that activation of basolateral amygdala neurons projecting to the nucleus accumbens core increases cue-induced approach to the location of reward, without influencing the pursuit of reward cues. Thus, these amygdala-to-accumbens neurons promote cue-induced expectation of reward, without changing the attractiveness of cues.

## INTRODUCTION

Through their predictive and conditioned reinforcing effects, reward-associated conditioned stimuli (CS) guide animals toward essential rewards such as water and food (Bindra, 1978). When a CS acquires predictive value, it evokes approach behaviors to the site of reward delivery, indicating conditioned anticipation of reward (Tolman, 1932; Hearst and Jenkins, 1974). When a CS also acquires conditioned reinforcing value, it can support the learning and performance of new instrumental actions, reinforced solely by the CS (Mackintosh, 1974; Robbins, 1978). Through their predictive and conditioned reinforcing properties, CS exert powerful control over psychology and behavior, promoting reward-seeking actions when the primary reward is not immediately available. However, when CS acquire too much incentive motivational value, they can promote pathological reward pursuit, such as in drug addiction (de Wit and Stewart, 1981; Vezina and Leyton, 2009). As such, gaining insight into the neural circuits mediating the effects of CS is relevant for understanding both adaptive and maladaptive appetitive behavior.

The psychological and behavioral effects of reward cues involve an overlapping network of several brain circuits (Cardinal et al., 2002; Saunders et al., 2018). In this context, the basolateral nucleus of the amygdala and its glutamatergic projections to the nucleus accumbens core [BLA→NAc core neurons (McDonald, 1996)] are a major focus of research (Everitt et al., 1999; Janak and Tye, 2015). BLA→NAc neurons preferentially fire in response to CS predicting rewarding rather than aversive stimuli (Beyeler et al., 2016). Pharmacological inhibition of the BLA also disrupts the ability of reward-predictive CS to increase both neuronal firing and dopamine release in the NAc core, two CS effects that mediate conditioned reward-seeking behavior (Ambroggi et al., 2008; Jones et al., 2010). Pharmacological disconnection of the BLA and NAc core also reduces CS-evoked instrumental responding for sucrose reward, suggesting that this projection is required for behavioral responding to appetitive cues (Ambroggi et al., 2008). However, because pharmacological manipulations are not circuit-specific, the preceding effects could also be due to an indirect action of BLA inactivation on NAc core activity. A circuit-specific, photoinhibition study suggests that neuronal activity in the BLA→NAc circuit is necessary for CS-triggered Pavlovian conditioned approach and consummatory responses (Stuber et al., 2011). However, this study did not distinguish between the core versus shell components of the NAc, and these can have different roles in cue-mediated behaviors (Ambroggi et al., 2011; West and Carelli, 2016).

Beyond triggering approach behaviors, reward-associated cues can also become conditioned reinforcers (Mackintosh, 1974; Robbins, 1978; Di Ciano and Everitt, 2004b), and it is unclear how activity in BLA→NAc core neurons might influence this effect. Decreasing the activity of BLA→NAc core neurons using optogenetics (Stefanik and Kalivas, 2013), chemogenetics (Puaud et al., 2021) or a pharmacological disconnection procedure (Di Ciano and Everitt, 2004a) disrupts CS-triggered increases in instrumental responding for cocaine reward when the drug is not available [but see (Everitt et al., 1989)]. However, the instrumental response was the same as that which previously led to cocaine. As such, these studies did not evaluate effects on the ability of a CS to support the learning of new instrumental actions, a critical test for conditioned reinforcement (Mackintosh, 1974; Robbins, 1978; Di Ciano and Everitt, 2004b). Other studies showed that when a CS reinforces the learning of new instrumental actions, excitotoxic lesions of the BLA do not influence the potentiation of instrumental responding produced by amphetamine infusions into the NAc core (Cador et al., 1989; Burns et al., 1993). This suggests that interactions between the BLA and the NAc core might not be required for the conditioned reinforcing properties of a CS. However, this has yet to be investigated using circuit-specific manipulations.

Thus, here we used *in vivo* optogenetic techniques to determine whether activating BLA→NAc core neurons influences (*i*) CS-triggered Pavlovian conditioned approach responses, and (*ii*) the ability of a CS to support the learning and performance of new instrumental actions.

## METHODS

### Animals

We used male Sprague-Dawley rats (225-275 g on arrival; Charles River Laboratories; from Montreal, Quebec, Canada, for Experiment 1; from Kingston, New York, United States, for Experiment 2, as the supplier changed locations). Rats were housed 3/cage for electrophysiological recordings or 1/cage for behavioral experiments, the latter to avoid damage to the optic implant from conspecifics. Rats were housed on a reverse dark-light cycle (lights off at 8:30 a.m.) and testing took place during the dark phase. Food and water were available *ad libitum*, except during behavioral testing where water access was restricted to 2 h/day to facilitate Pavlovian conditioning using water as the unconditioned stimulus (UCS; see ‘Pavlovian conditioning’ section). Experimental procedures involving rats were approved by the Université de Montréal’s ethics committee and followed the guidelines of the Canadian Council on Animal Care.

### Virus infusion and optic fiber implantation

Approximately 1 week after arrival to the animal colony, rats now weighing 325-350 g were anesthetised with isoflurane (5 % for induction, 2-3% for maintenance; CDMV, Saint-Hyacinthe, Quebec, Canada) and placed on a stereotaxic apparatus. At the beginning of the surgery, rats received a subcutaneous injection of carprofen (1.5 mg; CDMV) and an intramuscular injection of penicillin G procaine solution (3,000 IU; CDMV). The virus delivering ChR2-eYFP [AAV7-CamKIIa-hChR2(H134R)-eYFP; provided by Dr. Karl Deisseroth; Canadian Neurophotonics Platform Viral Vector Core Facility, RRID:SCR_016477, Quebec City, Quebec, Canada] was infused either bilaterally (Experiment 1) or unilaterally (Experiment 2; injected hemisphere was counterbalanced across groups) into the BLA (relative to Bregma: AP −2.8 mm, ML ±5.0 mm; relative to skull surface above the BLA: DV −8.2 mm). To this end, we used a Nanoject II (Drummond Scientific, PA, USA) coupled to a ~50-μm glass pipet to administer 27 microinjections of 36.8 nL each (23 nL/second, given every 10 seconds, total volume of ~1 μL/hemisphere). The glass pipet was left in place for 10 additional minutes after microinfusions. The craniectomy done to permit virus infusion was then sealed with bone wax (Ethicon, Somerville, New Jersey, United-States). For Experiment 2, control rats received a unilateral microinfusion into the BLA of an optically inactive virus (AAV7-CamKIIa-eYFP; Canadian Neurophotonics Platform Viral Vector Core Facility), using the same procedures. BLA projections to the NAc are mainly ipsilateral (Kita and Kitai, 1990; McDonald, 1996), and this ipsilateral connection is necessary for cue-triggered reward seeking (Ambroggi et al., 2008). Thus, for Experiment 2, an optic fiber (~300 μm core diameter, numerical aperture of 0.39, Thorlabs, Newton, New Jersey, United-States; glued with epoxy to a ferrule, model F10061F340, Fiber Instrument Sales Inc., Oriskany, New York, United-States) was implanted into the NAc core of the same hemisphere where virus was injected (relative to Bregma: AP +1.9 mm, ML ±1.6 mm; relative to skull surface: DV −6.7 mm). Then, six stainless steel screws were anchored to the skull and the optic implant was fixed in place with dental cement. Optogenetic stimulations occurred at least 3 weeks after the surgery, during which time rats were left in their home cages to recover.

### *In vivo* electrophysiology

We used *in vivo* electrophysiology to confirm laser-induced action potentials in BLA terminals expressing ChR2-eYFP in the NAc core. Anesthetized rats (urethane, 1.2 g/kg, i.p.) were placed inside a Faraday cage on a stereotaxic frame equipped with a body temperature controller. An optic fiber was implanted into the NAc core. This optic fiber was linked to the laser via a patch-cord built as described in Trujillo-Pisanty et al. (2015). Bone and dura above the BLA were removed and single-unit recordings were made using a glass micropipette filled with 2 M NaCl solution that contained 0.1% Fast green dye (impedance ranged from 2-6 mΩ at 1000 Hz). A silver wire (0.25 mm) inserted into the contralateral hemisphere served as a reference electrode. The micropipette was lowered with a hydraulic microdrive into the BLA to record single action potentials elicited by laser stimulation of ChR2-eYFP-expressing terminals in the NAc core (0-25 mW, 1-40 Hz, 5-ms pulse; 462-nm blue diode laser, Shanghai Laser & Optics Century Co. Ltd, Shanghai, China).

Antidromic action potentials elicited by laser stimulation of ChR2-eYFP-expressing terminals in the NAc core were fed into a high impedance headstage connected to a microelectrode amplifier (A-M Systems, Model 1800; Sequim, Washington, United-States). During laser stimulation, the low- and high-pass filters were set at 300 Hz and 5 kHz, respectively. Action potentials were displayed on an oscilloscope (Tektronix, Model TDS 1002; Beaverton, Oregon, United-States). The signal was digitalized and stored using DataWave recording (USB 16 channels) and DataWave SciWorks Experimenter Package (DataWave Technologies, Parsippany, New Jersey, United-States). At the end of the recording session, the recording site was marked by passing a 30-min, 25-μA direct cathodal current through the micropipette.

### Pavlovian conditioning

Behavioral training and testing took place in standard conditioning chambers (Med Associates, St. Albans, Vermont, United-States). During all sessions, a fan and a house-light were on unless otherwise stated. Water restriction began at least 9 days before the start of Pavlovian conditioning. Rats were habituated to water restriction by first receiving access to water bottles for 6 h/day for 4 days, then for 4 h/day for 3 days and finally for 2 h/day until the end of the experiment. Rats received at least 2 days of 2-h access/day prior to the start of Pavlovian conditioning, and water was always given at least 1 h after training/testing. As illustrated in Fig. 1A, rats were trained to associate a light-tone compound cue (illumination of cue lights for 5 s with concomitant extinction of house-light, followed by an 85-dB tone lasting 0.18 s) with delivery of 100 μL water (the UCS). CS-UCS pairings were presented 20 times per session, at a variable interval of 60 seconds. To determine whether the rats learned the CS-UCS association, we analysed approach to the water dish both during CS presentation [a measure of expectation of the primary reward (Tolman, 1932; Hearst and Jenkins, 1974)] and immediately after CS presentation, when water was delivered. To do so, in each conditioning session, we computed 4 measures; (1) the number of water dish visits during the 5-s CS minus the number of visits during the 5-s period preceding each CS presentation (pre-CS), (2) the number of CS trials with at least one dish visit, (3) the number of dish visits during the 5-s period following each CS presentation (post-CS) minus the number of visits during the 5-s pre-CS and (4) the latency to enter the water dish after CS presentation.

**Figure 1.**
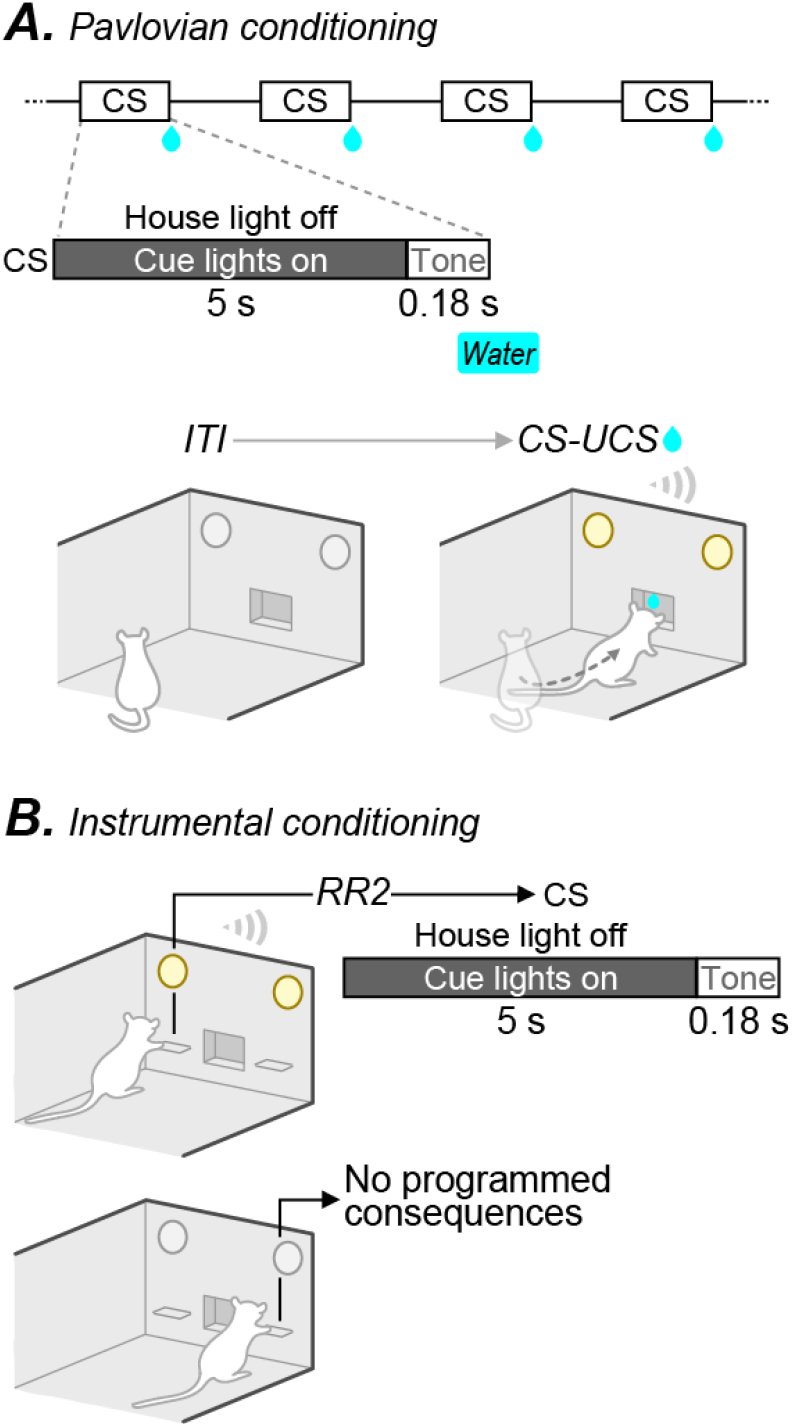
Pavlovian and instrumental conditioning. (**A**) During Pavlovian conditioning, water-restricted rats learned that a light-tone cue (CS) predicts water delivery (100 μL; UCS, 20 CS-UCS pairings per session). (**B**) After learning the CS-UCS contingency, rats were given access to two new levers. Pressing on the active lever led to CS presentation alone (RR2 schedule; no water), whereas pressing on the inactive lever had no programmed consequences.

### Conditioned reinforcement test

After Pavlovian conditioning, we evaluated the conditioned reinforcing effects of the CS. To this end, we examined whether rats would spontaneously learn to press a lever for CS presentations, without water delivery (Mackintosh, 1974; Robbins, 1978). This procedure dissociates incentive motivation for the CS versus UCS, because lever responding is new to the rats, and was never previously reinforced by the UCS. Rats were given instrumental conditioning sessions, during which they were presented with two levers they had not seen before. As Fig. 1B illustrates, presses on one (active) lever produced the CS under a random-ratio 2 schedule (RR2). Presses on the active lever during CS presentation or on the other (inactive) lever were recorded but had no programmed consequences. Rats were first allowed to spontaneously learn to lever press for the CS during instrumental conditioning sessions. These sessions ended after 10 presses on the active lever or after 30 minutes. Rats received one session per day until they earned at least one CS presentation (this took 1-2 sessions for all rats). The day after completion of this training, we measured instrumental responding for the CS in conditioned reinforcement test sessions. These sessions were like instrumental conditioning sessions, except that they had a fixed duration of 20 minutes, and no maximum number of active lever presses.

### Experiment 1: Effects of optogenetic stimulation on action potentials in BLA→NAc core neurons expressing ChR2-eYFP

As shown in Fig. 2A, rats (n = 2) received the ChR2-eYFP virus into the BLA. Three weeks later, rats were anesthetised using urethane (1.2 g/kg, i.p.) and antidromic action potentials were recorded from BLA neurons following laser application via an optic fiber in the ipsilateral NAc core.

**Figure 2.**
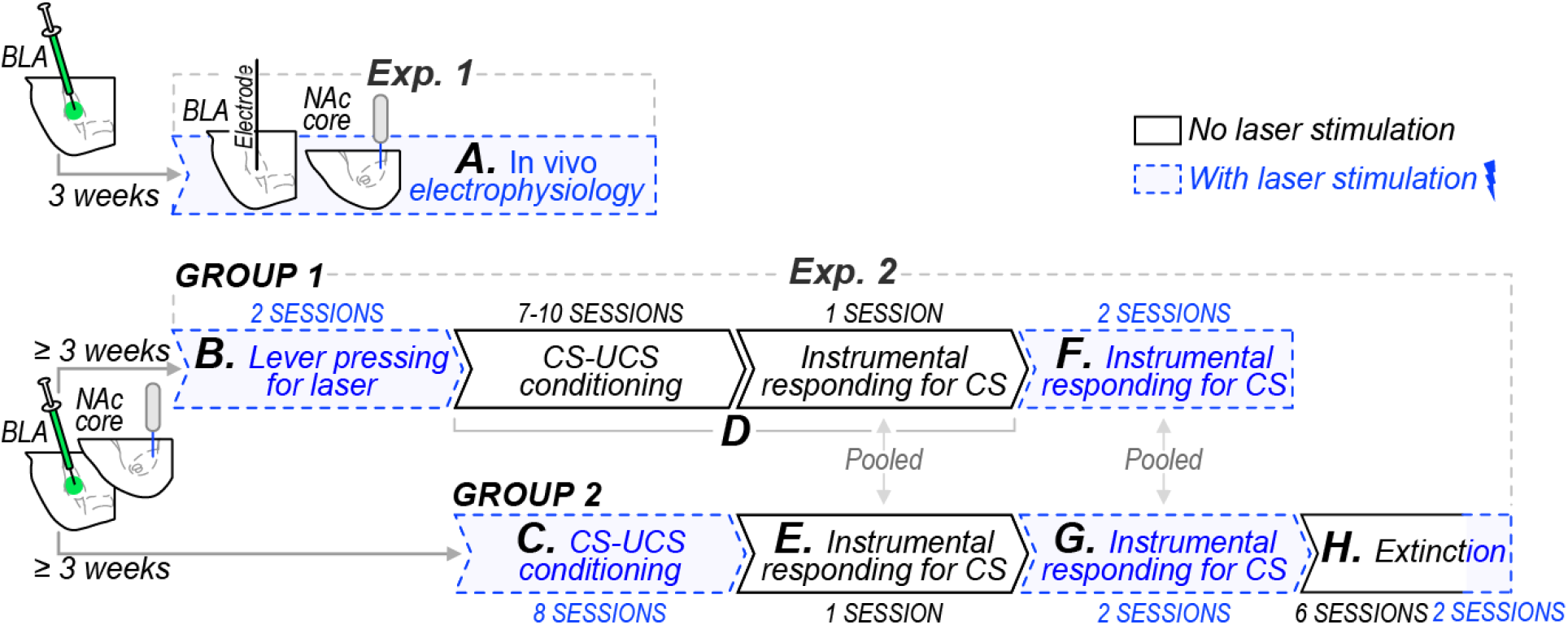
Experimental timelines. (**A**) In Experiment 1, rats received AAV7-CamKIIa-hChR2(H134R)-eYFP into the BLA of both hemispheres. Three weeks later, we recorded action potentials in the BLA evoked by laser stimulation in the NAc core using *in vivo* electrophysiology. (**B**-**H**) In Experiment 2, two groups of rats received the ChR2-eYFP virus (AAV7-CamKIIa-hChR2(H134R)-eYFP) or the optically inactive virus delivering eYFP alone (AAV2-CamKIIa-eYFP) in the BLA of one hemisphere, and an optic fiber was chronically implanted into the NAc core of the same hemisphere. At least 3 weeks later, we assessed the effects of optogenetic stimulation of BLA→NAc core neurons on (**B**) lever pressing behavior, (**C**) CS-UCS conditioning, (**D**-**G**) instrumental responding for the CS, and (**H**) extinction.

### Experiment 2

Two independent groups of rats were used in Experiment 2. Figs. 2B-H illustrate the experimental timeline for each group.

#### Effects of optogenetic stimulation of BLA→NAc core neurons on lever pressing behavior

Here we determined whether optogenetic stimulation of BLA→NAc core neurons is intrinsically rewarding and would therefore reinforce lever pressing behavior. This was important to address, because if rats lever press for optogenetic stimulation alone, this would confound interpretation of results from subsequent experiments on conditioned reinforcement. At least three weeks after virus injection, ChR2-eYFP rats (n = 8) and eYFP rats (n = 5) could lever press for laser stimulation (20-Hz, 5-ms pulse, 10 mW, 5.18-s duration) combined with presentation of the light-tone compound stimulus under a fixed-ratio schedule of 1 (FR1; Fig. 2B). This was the same light-tone stimulus later used for Pavlovian conditioning, except that at this stage, it was new to the rats, and it had not previously been paired with water or any other outcome. Rats received 2 sessions lasting 30 min each (1 session/day, on 2 consecutive days). Because there were no significant differences in lever pressing behavior and in the number of stimulations earned between the sessions, we pooled data across sessions in the final analysis. After these lever-pressing tests, one ChR2-eYFP rat lost his headcap and was eliminated from subsequent testing (described under ‘*Effects of optogenetic stimulation of BLA→NAc core neurons during Pavlovian conditioning on later instrumental responding for the CS’)*.

#### Effects of optogenetic stimulation of BLA→NAc core neurons on the development of CS-triggered conditioned approach behavior

Here we determined whether optogenetic stimulation of BLA→NAc core neurons during Pavlovian conditioning enhances the acquisition of CS-triggered conditioned approach to the site of water delivery, a measure of CS-induced anticipation of reward. At least 3 weeks after virus injection, 3 groups were formed (ChR2-eYFP/laser, n = 11; ChR2-eYFP/no laser, n = 6; and eYFP-laser n = 8), and all groups received 8 Pavlovian conditioning sessions (1 session/day, on consecutive days; Fig. 2C, Group 2). During conditioning, rats in the ChR2-eYFP/laser and eYFP/laser groups received CS-paired laser stimulation (20-Hz, 5-ms pulses, 10 mW, 5.18-s duration).

#### Effects of optogenetic stimulation of BLA→NAc core neurons during Pavlovian conditioning on later instrumental responding for the CS

We determined whether optogenetic stimulation of BLA→NAc core neurons during prior Pavlovian conditioning influences lever pressing for the CS, a measure of the CS’s conditioned reinforcing properties. To evaluate this, we used rats from groups 1 and 2. As Fig. 2D shows, rats from group 1 received 7-10 Pavlovian conditioning sessions without laser stimulations (ChR2-eYFP/no laser, n = 7; eYFP, n = 5). As described above (see also Fig. 2C), rats from group 2 received 8 Pavlovian conditioning sessions, with or without laser stimulation (ChR2-eYFP/laser, n = 11; ChR2-eYFP/no laser, n = 6; eYFP-laser, n = 8). Following this, all rats from both groups were tested for conditioned reinforcement, as described under ‘*Conditioned reinforcement test*’. Briefly, rats could lever press for CS presentations, without laser stimulation, as Figs. 2D-E show.

#### Effects of optogenetic stimulation of BLA→NAc core neurons during instrumental responding for a CS

In rats previously naïve to optogenetic stimulation, we determined whether CS-paired laser stimulations influence the conditioned reinforcing value of the CS. To this end, we used the rats that had received Pavlovian conditioning without optogenetic stimulation (ChR2-eYFP/no laser during Pavlovian conditioning: n = 7 from group 1, n = 6 from group 2; eYFP: n = 5 from group 1, n = 8 from group 2). As Figs. 2F-G show, these rats now received 2 conditioned reinforcement tests with laser stimulation. On the first test, each CS presentation was paired with laser stimulation (20-Hz, 5-ms pulses, 10 mW, 5.18-s duration). On the second test, laser stimulation was explicitly unpaired from CS presentation to assess any non-specific effects of optogenetic stimulation on instrumental responding. Thus, laser stimulation was given 3 seconds after each CS presentation (20-Hz, 5-ms pulses, 10 mW, 5.18-s duration).

#### Effects of optogenetic stimulation of BLA→NAc core neurons on extinction responding

When the CS no longer predicts the UCS, extinction learning occurs, whereby CS-triggered responses (Pavlovian or Instrumental) progressively abate. Extinction learning is thought to involve the formation of new CS-no UCS associations that inhibit initially learned CS-UCS associations (Rescorla, 1993). Our objectives here were to determine (1) how optogenetic stimulation of BLA→NAc core neurons during initial Pavlovian conditioning influences later extinction of Pavlovian conditioned approach behavior, and (2) how optogenetic stimulation of BLA→NAc core neurons *during* extinction sessions influences Pavlovian conditioned approach behavior. As Fig. 2H shows, we used rats from group 2 to address these questions. Following conditioned reinforcement tests, the rats received 2 reminder CS-UCS conditioning sessions (without laser) and then 2 extinction sessions during which the CS was presented, but no water was delivered (without laser). Because these rats had received their initial Pavlovian conditioning with laser stimulation (Fig. 2C), this allowed us to determine how optogenetic stimulation of BLA→NAc core neurons during previous conditioning might influence later extinction of conditioned approach responses. Furthermore, to determine whether optogenetic stimulation of BLA→NAc core neurons during extinction sessions influences Pavlovian conditioned approach behavior, the same rats received 2 more reminder CS-UCS conditioning sessions, during which each CS presentation was followed by water delivery (without laser). The rats then received 2 extinction sessions during which each CS presentation was now paired with laser stimulation (20-Hz, 5-ms pulses, 10 mW, 5.18-s duration).

### Histology and immunohistochemistry

Anesthetised rats (urethane, 1.2 g/kg, i.p.) were intracardially perfused first with phosphate buffered saline (PBS) and then with 4 % paraformaldehyde. Brains were extracted, post-fixed in 4 % paraformaldehyde for 1 hour, then stored in 30 % sucrose/PBS for 23 hours. Brains were stored at −20 °C and then 40-μm coronal slices were sectioned in a cryostat and stored in antifreeze solution at −20 °C until processing. For immunohistochemistry, all steps were done with gentle agitation and at room temperature unless otherwise stated. Free-floating slices were washed 3 times for 15 minutes in PBS, post-fixed in 4% paraformaldehyde for 30 minutes and washed again 3 times for 15 minutes each time in PBS. Slices were then incubated for 60 minutes in blocking solution containing 0.3 % triton X-100, 10 % bovine serum albumin (BSA), 0.02 % sodium azide and 5 % goat serum. Following this, slices were incubated in the same blocking solution as above but now containing 0.5 % BSA and 1:2000 of a chicken polyclonal antibody against GFP (AvesLabs, product # GFP-1020; Cedarlane, Burlington, Ontario, Canada) for 24 hours at 4° C. The next day, slices were washed 3 times for 15 minutes in PBS. Slices were then incubated in the blocking/0.5 % BSA solution containing 1:500 of a secondary antibody (goat anti-chicken, Alexa Fluor 488, Invitrogen, # A-11039; ThermoFisher Scientific, Burlington, Ontario, Canada) for 2 hours in a dark room. Slices were washed again 3 times for 15 minutes each time in PBS and were then fixed on microscope slides with mounting solution and coverslips. ChR2-eYFP expression was visualised with a fluorescent microscope and optic fiber placement was estimated based on the Paxinos and Watson (1986) atlas.

### Statistics

During laser self-stimulation tests, we used an unpaired *t*-test to analyse group differences in self-administered laser stimulations, and mixed-model ANOVA to analyse group differences in lever pressing behavior (Group × Lever type: ‘Lever type’ as a within-subjects variable). To analyse effects of laser stimulation on conditioned approach responses during Pavlovian conditioning, we used mixed-model ANOVA to compare the groups on water dish visits during and after CS presentation, number of CS trials with a response, and latency to enter the water dish after the CS, across sessions (Group × Session, ‘Session’ as a within-subjects variable). To analyse effects of laser stimulation on instrumental responding for conditioned reinforcement, we used mixed-model ANOVA to compare the groups on lever pressing for the CS (Group × Lever type, ‘Lever type’ as a within-subjects variable). We used either one-way ANOVA (Fig. 6C) or unpaired *t*-tests (Figs. 7C and 7E) to analyse group differences in the number of CS presentations earned. To assess effects of laser stimulation on behavior during extinction sessions, we used mixed-model ANOVA to compare the groups on water dish visits during and after CS presentation, number of CS trials with a response and latency to enter the water dish after CS presentation, across sessions or trials (Group × Session or Trial, ‘Session’ and ‘Trial’ as within-subjects variables). When interaction and/or main effects were significant (*p* < 0.05), effects were analysed further using Bonferroni-adjusted multiple post-hoc comparisons. Values in figures are means ± SEM.

## RESULTS

### Experiment 1: Effects of optogenetic stimulation on action potentials in BLA→NAc core neurons expressing ChR2-eYFP

Fig. 3A illustrates representative recordings of antidromic actions potentials in the BLA elicited by laser stimulation in the NAc core of rats that had received the ChR2-eYFP virus in the BLA 3 weeks before. Under our conditions, laser stimulation produced reliable action potentials in ChR2-eYFP-expressing BLA→NAc core neurons.

**Figure 3.**
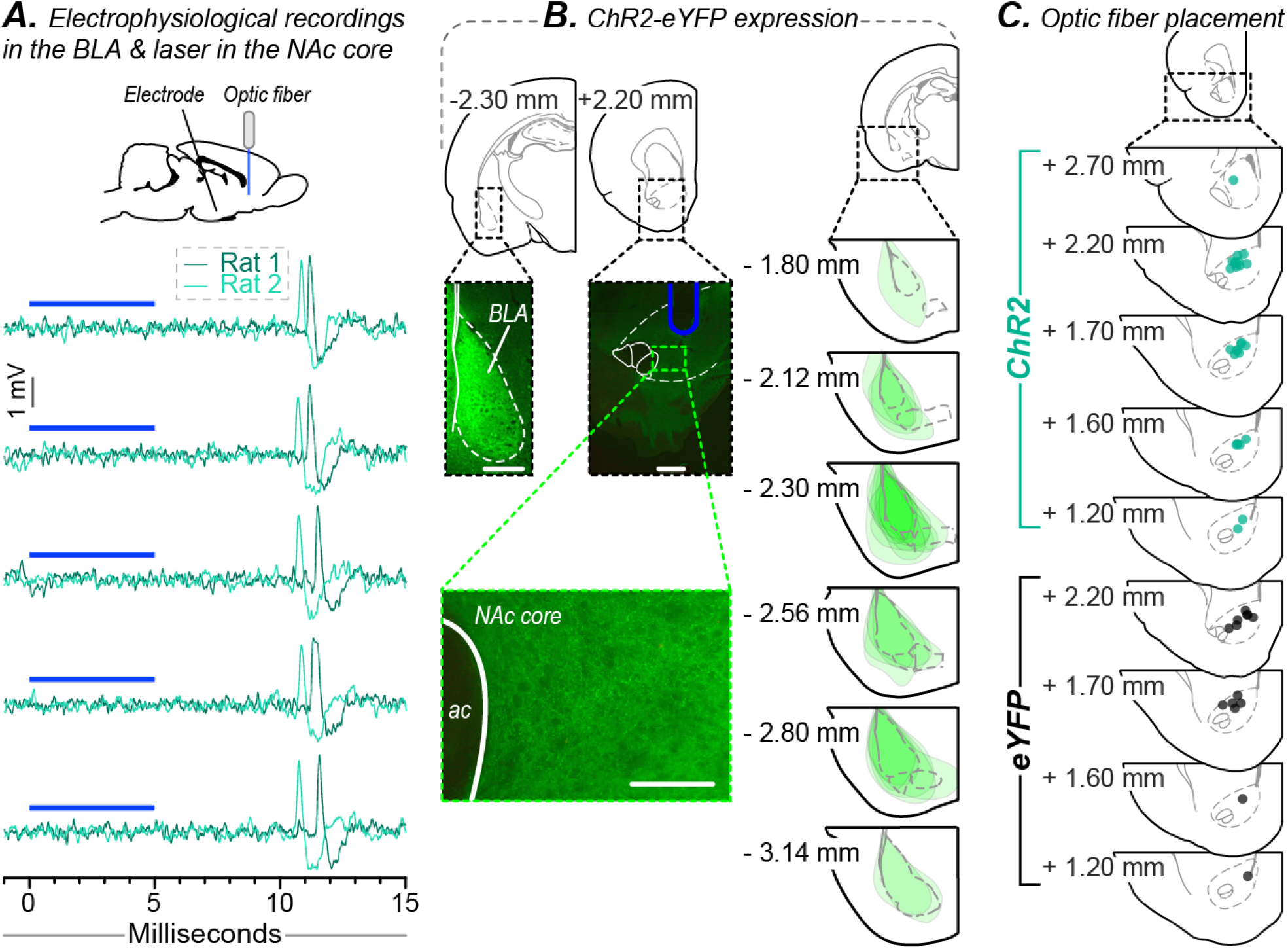
Electrophysiological recordings of laser-induced actions potential in BLA→NAc core neurons, ChR2-eYFP expression and optic fiber placement. (**A**) Representative electrophysiological recordings in the BLA of antidromic action potentials elicited by laser application in the NAc core 3 weeks after virus injection into the BLA. (**B**) On the left, pictures of representative ChR2-eYFP expression in the BLA and in BLA terminals in the NAc core (scale bars: 1 mm in top 2 pictures and 50 μm in bottom picture). On the right, schematic representation of ChR2-eYFP expression in the BLA at the anteroposterior level with the highest density of viral expression, for each ChR2-eYFP rat from Experiment 2. (**C**) Estimated placement of optic fibers in the NAc core of ChR2-eYFP (turquoise dots) and eYFP (black dots) rats from Experiment 2.

### Experiment 2

Fig. 3B (left panels) shows representative brain sections illustrating the expression of ChR2-eYFP in the BLA and in BLA terminals in the NAc core. Fig. 3B (right panels) also shows a schematic representation of viral expression in the BLA at the anteroposterior level with the highest ChR2-eYFP density, for each ChR2-eYFP rat used in Experiment 2. Fig. 3C shows estimated placement of optic fibers in the NAc core of the ChR2-eYFP and eYFP rats used in Experiment 2. All rats had optic fibers placed in the NAc core.

### Effects of optogenetic stimulation of BLA→NAc core neurons on lever pressing behavior

Here we determined whether optogenetic stimulation of BLA→NAc core neurons supports lever pressing behavior, and we found that it did not. As Fig. 4A shows, pressing on the active lever produced optogenetic stimulation combined with presentation of a light-tone stimulus, under a FR1 schedule of reinforcement. Pressing on the inactive lever had no programmed consequences. Fig. 4B shows that across ChR2-eYFP and eYFP groups, rats pressed more on the active versus inactive lever (effect of Lever Type, *F*_(1,11)_ = 15.89, *p* = 0.002). This suggests that the light-tone stimulus has at least some intrinsic reinforcing properties [see also (Kish, 1954; Stewart and Hurwitz, 1958)]. Importantly, there were no group differences in lever pressing (Group × Lever type interaction or Group effects; all *P*’s > 0.05), indicating that optogenetic stimulation of BLA→NAc core neurons does not reinforce operant behavior. In further support, ChR2-eYFP and eYFP rats earned a similar number of laser stimulations (Fig. 4C; *p* > 0.05). Thus, rats do not reliably self-administer optogenetic stimulation of BLA→NAc core neurons, indicating that this stimulation is not reinforcing.

**Figure 4.**
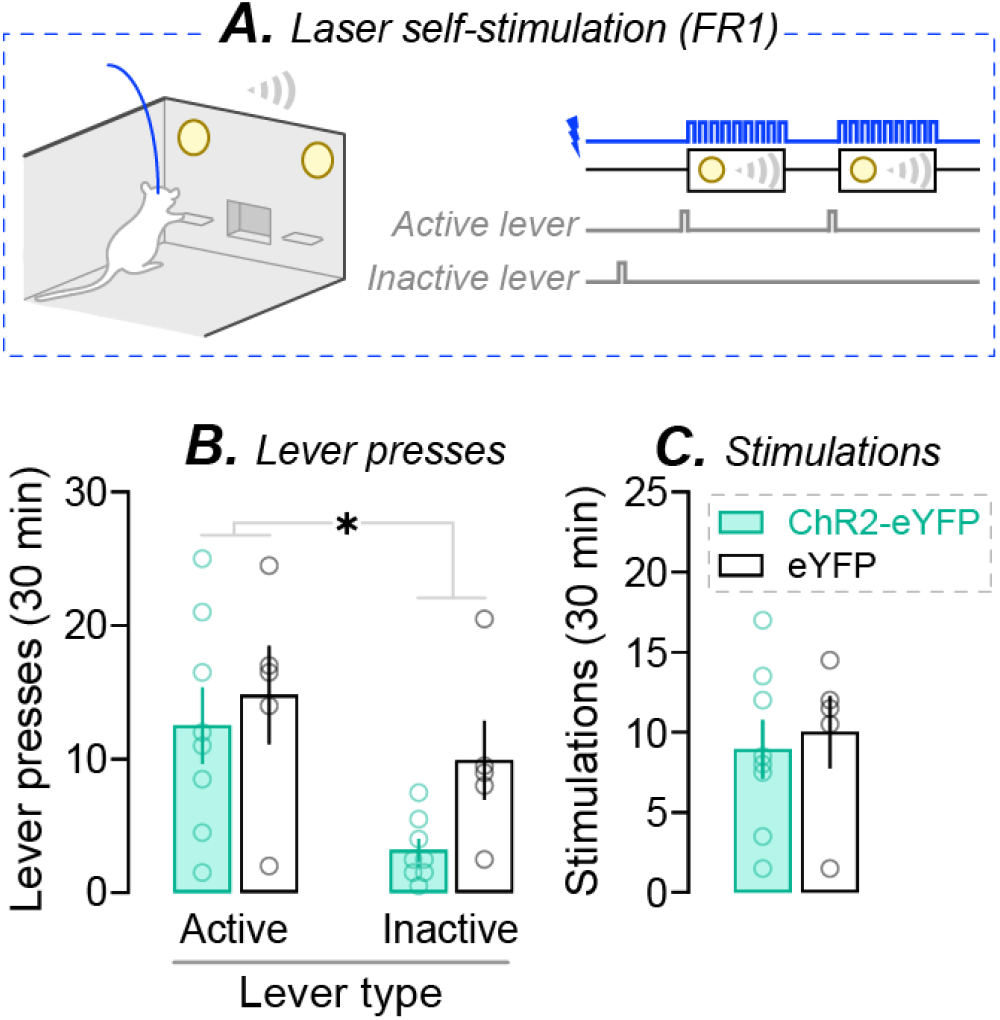
Optogenetic stimulation of BLA→NAc core neurons does not reinforce instrumental responding. (**A**) Rats were free to press on two levers. Pressing on the active lever produced optogenetic stimulation of BLA→NAc core neurons paired with presentation of a light-tone cue, under a FR1 schedule. Pressing on the inactive lever had no programmed consequences. ChR2-eYFP and eYFP rats (**B**) pressed more on the active versus inactive lever, with no group differences, and (**C**) self-administered a similar number of stimulations. eYFP rats, n = 5; ChR2-eYFP rats, n = 8. In (**B**); \**p* < 0.05, main effect of Lever type. Values in figures are means ± SEM. Individual data are shown on histograms.

### Effects of optogenetic stimulation of BLA→NAc core neurons on the development of CS-triggered conditioned approach behavior

Here we determined the effects of optogenetic stimulation of BLA→NAc core neurons on the acquisition of CS-evoked conditioned approach behavior (Fig. 5A). During each CS-UCS conditioning session, we analysed four measures of CS-triggered conditioned approach behavior: (1) the number of dish visits during each 5-s CS minus visits during the 5-s period preceding each CS presentation (pre-CS), (2) the number of CS trials with at least one dish visit during the CS, (3) the number of water dish visits during the 5-s period following each CS presentation (post-CS) minus visits during the pre-CS, and finally, (4) the latency to enter the water dish after CS presentation. Control rats included ChR2-eYFP rats that received no laser stimulation (n = 6) and eYFP rats that received CS-paired laser stimulation during Pavlovian conditioning (n = 8). There were no behavioral differences between these 2 groups, and they were pooled (Controls, n = 14).

**Figure 5.**
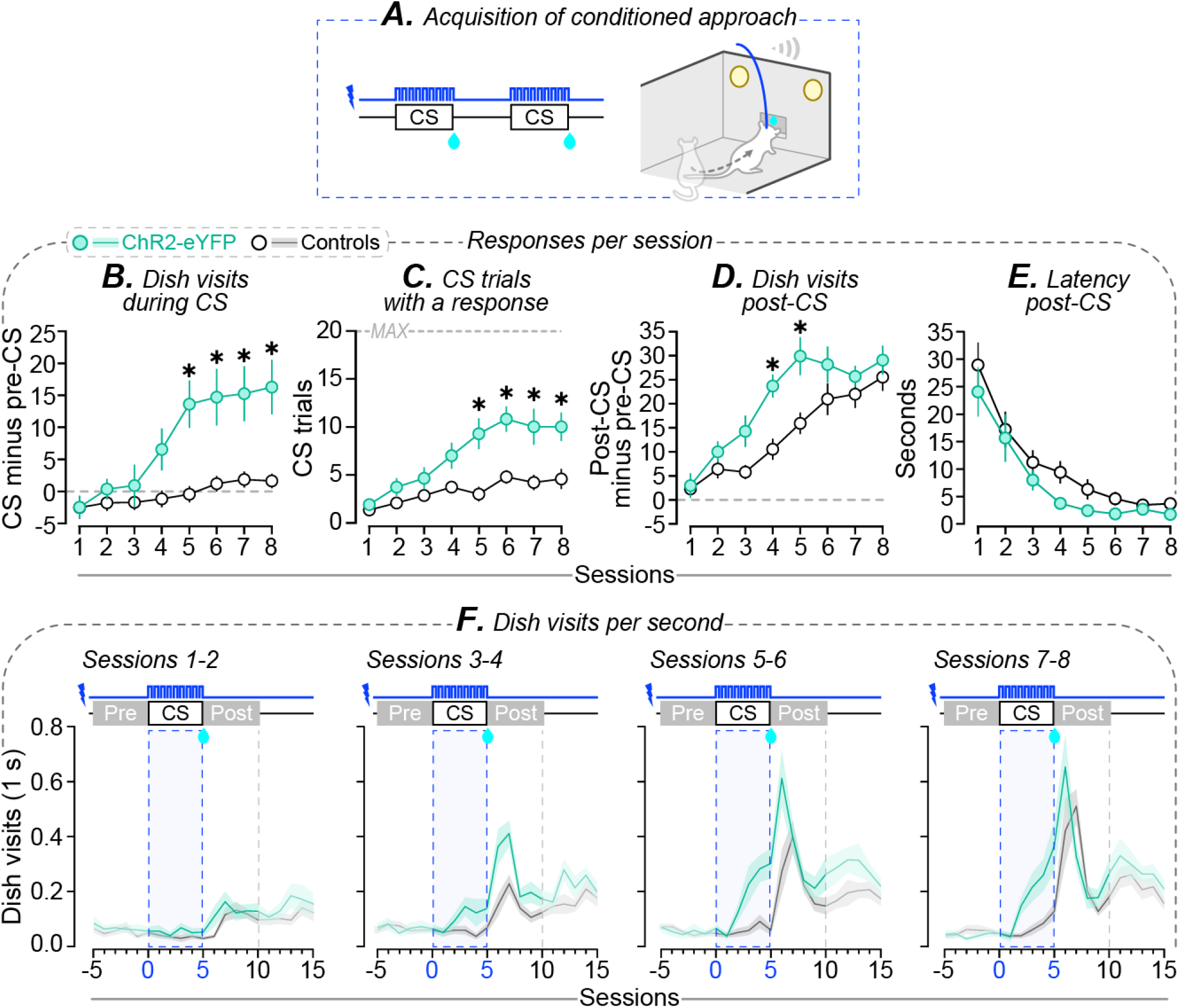
Optogenetic stimulation of BLA→NAc core neurons promotes the acquisition of CS-evoked conditioned approach behavior. (**A**) Water-restricted rats received Pavlovian conditioning sessions where they learned that a light-tone CS predicts water delivery. CS presentation was combined with optogenetic stimulation of BLA→NAc core neurons. The control group included ChR2-eYFP rats that did not receive laser stimulation and eYFP rats that did. Compared to controls, ChR2-eYFP rats showed greater increases in (**B**) the number of dish visits during the 5-s CS minus the number of dish visits in the 5 s prior to the CS (pre-CS), (**C**) the number of CS trials with at least one dish visit during the CS, and (**D**) the number of dish visits 5 s after each CS minus the number of visits during the pre-CS. (**E**) The latency to enter the water dish after CS presentation progressively decreased across sessions, and this was similar between the ChR2-eYFP and control groups. These results indicate that all rats learned the CS-UCS contingency, and that ChR2-eYFP rats showed potentiated CS-triggered conditioned approach relative to controls. (**F**) Dish visits per second before, during and after CS presentation, averaged over sessions 1-2, 3-4, 5-6 and 7-8. Control rats, n = 14; ChR2-eYFP rats, n = 11. In (**B**-**D**); \**p* < 0.05, relative to controls in the same session. Values in figures are means ± SEM.

#### Pairing optogenetic stimulation of BLA→NAc core neurons with CS presentation enhanced CS-evoked conditioned approach behavior

ChR2-eYFP rats visited the water dish more often *during* CS presentations than controls did, and did so from Session 5 onwards (Fig. 5B; Group × Session interaction, *F*_(7,161)_ = 6.06, *p* < 0.0001; Group effect, *F*_(1,23)_ = 12.32, *p* = 0.002; ChR2-eYFP rats > controls; Session 5, *p* = 0.0004; Session 6, *p* = 0.0007; Session 7, *p* = 0.0008; Session 8, *p* = 0.0002). Furthermore, across sessions, only ChR2-eYFP rats significantly increased their rate of water dish visits during CS presentations (Fig. 5B; Session effect, *F*_(7,161)_ = 14.02, *p* < 0.0001; ChR2-eYFP rats; relative to Session 1: < Session 4, *p* = 0.0058; < Session 5, *p* < 0.0001; < Session 6, *p* < 0.0001; < Session 7, *p* < 0.0001; < Session 8, *p* < 0.0001). In contrast, control rats maintained a low and relatively stable rate of water dish visits during CS presentations (all *P*’s > 0.05). Across groups, rats increased the number of CS trials during which they visited the water dish across sessions (Fig. 5C; Session effect, *F*_(7,161)_ = 17.75, *p* < 0.0001). However, this increase was more pronounced in ChR2-eYFP rats relative to controls (Fig. 5C; Group × Session interaction, *F*_(7,161)_ = 4.61, *p* = 0.0001; Group effect, *F*_(1, 23)_ = 14.62, *p* = 0.0009; ChR2-eYFP rats > controls: Session 5, *p* = 0.0001; Session 6, *p* = 0.0003; Session 7, *p* = 0.0006; Session 8, *p* = 0.002) and also occurred earlier in ChR2-eYFP rats (relative to Session 1; ChR2-eYFP rats: < Session 4, *p* < 0.0001; < Session 5, *p* < 0.0001; < Session 6, *p* < 0.0001; < Session 7, *p* < 0.0001; < Session 8, *p* < 0.0001; controls: < Session 6, *p* = 0.0058; < Session 7, *p* = 0.04; < Session 8, *p* = 0.01). Thus, both ChR2-eYFP and control rats learned the CS-UCS contingency, but the ChR2-eYFP rats showed enhanced CS-triggered conditioned approach behavior, and did so following fewer conditioning sessions.

There were also group differences in conditioned approach responses emitted immediately following CS presentation (‘post-CS’ responses). Across sessions, both ChR2-eYFP and control rats increased their number of post-CS water dish visits (Fig. 5D; Session effect, *F*_(7,161)_ = 36.37, *p* < 0.0001). ChR2-eYFP rats showed accelerated acquisition of this response, such that on Sessions 4-5, they showed more post-CS water dish visits compared to control rats (Fig. 5D; Group × Session interaction, *F*_(7,161)_ = 2.56, *p* = 0.02; Group effect, *F*_(1,23)_ = 7.34, *p* = 0.01; ChR2-eYFP rats > controls; Session 4, *p* = 0.0049; Session 5, *p* = 0.002. No other comparisons were statistically significant). Both ChR2-eYFP and control rats also progressively decreased their latency to enter the water dish after CS presentation (Fig. 5E; Session effect, *F*_(7,161)_ = 41.37, *p* < 0.0001), and there were no group differences in this response (all *P*’s > 0.05). Thus, relative to controls, ChR2-eYFP rats showed enhanced conditioned approach behaviors immediately following CS presentation, and also acquired this response earlier during conditioning.

#### Pairing optogenetic stimulation of BLA→NAc core neurons with CS presentation increased approach behavior specifically during CS-UCS trials, without producing a general increase in approach behavior to the site of water delivery

When we excluded water dish visits made during CS presentations or during the post-CS period, group differences in water dish entries disappeared (data not shown; Group × Session interaction or Group effects; all *P*’s > 0.05). Fig. 5F further highlights this effect. The figure illustrates dish visits per second before, during and after CS presentation, and shows that optogenetic stimulation of BLA→NAc core neurons progressively and selectively increased CS-triggered conditioned approach behavior to the site of reward delivery.

In summary, both ChR2-eYFP rats and controls acquired the CS-UCS association across conditioning sessions, as all rats made more, and increasingly more rapid visits to the water dish after CS presentation. However, compared to control rats, ChR2-eYFP rats showed more conditioned approach behavior *during* CS presentations (Figs. 5B-C), and they also required fewer training sessions to show a robust conditioned approach response after CS presentation (Fig. 5D). These findings suggest that optogenetic stimulation of BLA→NAc core neurons potentiates CS-triggered conditioned approach responses.

### Effects of optogenetic stimulation of BLA→NAc core neurons during Pavlovian conditioning on later instrumental responding for the CS

Here we determined whether having received BLA→NAc core neuron stimulation during prior CS-UCS conditioning changes the incentive motivational properties of that CS, as measured by the spontaneous acquisition of a lever-pressing response reinforced solely by the CS (*i.e*., instrumental responding for conditioned reinforcement). We examined this in rats from groups 1 (Fig. 2D) and 2 (Fig. 2E). Rats from group 2 were ChR2-eYFP and eYFP rats that had already received Pavlovian conditioning (Fig. 5). Rats from group 1 (ChR2-eYFP and eYFP rats) had previously been used to determine whether photo-stimulations of BLA→NAc core neurons reinforce lever pressing behavior (Fig. 4). These rats now received 7-10 Pavlovian conditioning sessions without laser stimulation, during which they learned to associate a light-tone CS with a water UCS. ChR2-eYFP and eYFP rats from group 1 showed similar acquisition of the CS-UCS contingency, as indicated by (*i*) a comparable across-session increase in dish visits made both during and immediately after CS presentation (data not shown; Session effect; dish visits during CS minus dish visits during pre-CS, *F*_(6, 60)_ = 9.96, *p* < 0.0001; post-CS dish visits minus pre-CS dish visits, *F*_(6,60)_ = 17.03, *p* < 0.0001; Group and Group × Session interaction effects, all *P*’s > 0.05), (*ii*) a comparable increase in the number of CS trials during which rats visited the water dish at least once (Session effect, *F*_(6, 60)_ = 15.72, *p* < 0.0001; Group and Group × Session interaction effects, all *P*’s > 0.05) and, (*iii*) a comparable decrease in the latency to enter the water dish after CS presentation (Session effect, *F*_(6,60)_ = 8.66, *p* < 0.0001; Group and Group × Session interaction effects, all *P*’s > 0.05). Thus, all rats learned the CS-UCS contingency prior to conditioned reinforcement tests, and the findings also indicate that mere expression of ChR2-eYFP in BLA neurons does not influence Pavlovian conditioning to a water-paired cue, in accordance with previous work (Servonnet et al., 2020).

Following Pavlovian CS-UCS conditioning, rats received a single session where they could press a lever that produced the CS alone (no water and no laser stimulation) or an inactive lever that had no programmed consequences (Fig. 6A). During this session, rats from groups 1 and 2 showed similar rates of lever pressing behavior. Thus, ChR2-eYFP rats from group 1 were pooled with ChR2-eYFP/no laser rats from group 2 (‘ChR2-eYFP/no laser during Pavlovian conditioning’, n = 13), and eYFP rats were also pooled across the 2 groups (n = 13).

**Figure 6.**
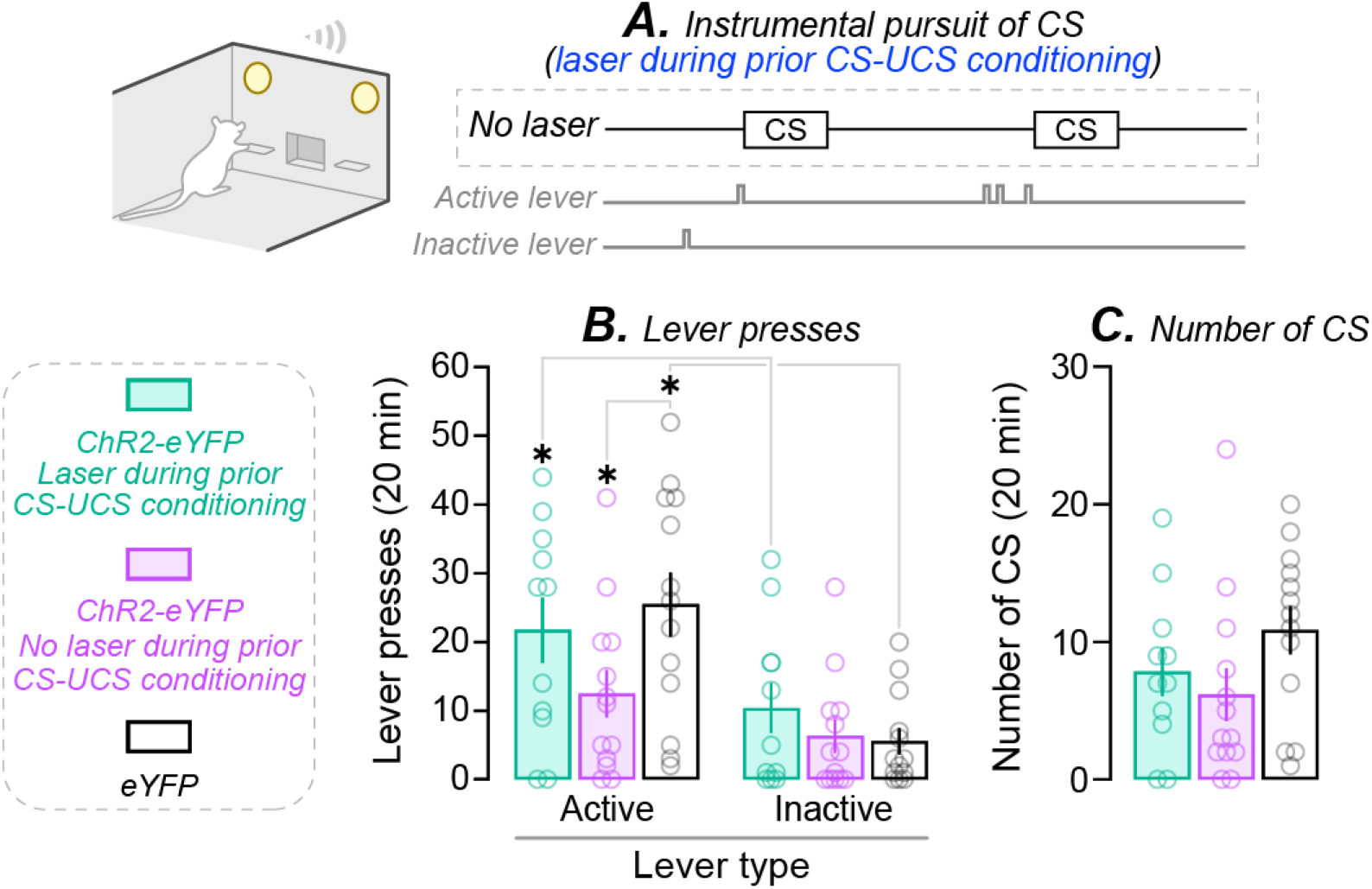
Optogenetic stimulation of BLA→NAc core neurons during prior CS-UCS conditioning does not subsequently alter instrumental responding for CS. (**A**) We examined the effects of stimulating BLA→NAc core neurons during Pavlovian conditioning on subsequent lever-pressing for the CS alone, without the water UCS. During initial CS-UCS conditioning sessions, ChR2-eYFP rats received CS presentation combined with optogenetic stimulations (turquoise histograms). Control rats received CS-UCS conditioning without optogenetic stimulation (ChR2-eYFP, purple histograms; eYFP, white histograms). Following this, rats received a session where they could lever press on an active lever for CS presentations (RR2 schedule) and on an inactive lever. (**B**) ChR2-eYFP rats that had received prior Pavlovian conditioning with optogenetic stimulation and eYFP rats pressed more on the active lever relative to the inactive lever, indicating that the CS acquired conditioned reinforcing value. The ChR2-eYFP rats that had received prior CS-UCS conditioning without laser stimulation did not discriminate between the levers. (**C**) Rats across groups earned a similar number of CS presentation during the conditioned reinforcement test. ChR2-eYFP rats that had received prior CS-UCS conditioning with optogenetic stimulation, n = 11; ChR2-eYFP rats that had received prior CS-UCS conditioning without optogenetic stimulation, n = 13; eYFP rats, n = 13. In (**B**); \**p* < 0.05. Values in figures are means ± SEM. Individual data are shown on histograms.

#### Optogenetic stimulation of BLA→NAc core neurons during prior Pavlovian conditioning does not change later instrumental responding for the CS

Fig. 6B shows lever pressing behavior of: ChR2-eYFP rats that had received prior CS-UCS conditioning with (turquoise) or without (purple) CS-paired optogenetic stimulations of BLA→NAc core neurons and eYFP rats (white). Across groups, rats pressed more on the active versus inactive lever (Fig. 6B; Lever effect, *F*_(1,34)_ = 33.33, *p* < 0.0001). Thus, rats spontaneously acquired and performed a new lever-pressing response to obtain the CS alone, indicating that the CS had become a conditioned reinforcer. This effect also depended on the group (Group × Lever type interaction, *F*_(2,34)_ = 3.65, *p* = 0.04). Further analysis of this interaction effect showed that some groups significantly distinguished between the active versus inactive lever, while others did not. Specifically, only eYFP control rats and ChR2-eYFP rats that had received optogenetic stimulation during prior CS-UCS conditioning pressed more on the active relative to the inactive lever (Fig. 6B; active > inactive: white histograms, *p* < 0.0001; turquoise histograms, *p* = 0.02). ChR2-eYFP rats that had received prior CS-UCS conditioning without laser stimulation did not significantly discriminate between the levers (Fig. 6B; purple histograms, *p* > 0.05). These rats also pressed less on the active lever compared to eYFP rats (Fig. 6B; active lever; white > purple, *p* = 0.03. No other comparisons were significant). Importantly, ChR2-eYFP rats that had received laser stimulations during prior CS-UCS conditioning and eYFP control rats showed similar lever pressing behavior during the conditioned reinforcement test. This suggests that optogenetic stimulation during prior CS-UCS conditioning does not change the conditioned reinforcing properties of the CS. In further support of this, the three groups earned a similar number of CS presentations during the conditioned reinforcement test (Fig. 6C; Group effect, *p* > 0.05).

Thus, pairing optogenetic stimulation of BLA→NAc core neurons with CS presentations during Pavlovian conditioning does not influence the operant pursuit of that CS. This suggests that activation of BLA→NAc core neurons when rats are learning a CS-UCS contingency does not alter the attribution of conditioned reinforcing value to that CS.

### Effects of optogenetic stimulation of BLA→NAc core neurons during instrumental responding for a CS

In the previous test, we examined how stimulating BLA→NAc core neurons during initial CS-UCS conditioning influences later instrumental responding for the CS. Here we examined the effects of stimulating these neurons during instrumental responding for the CS (Fig. 7A). To this end, we used ChR2-eYFP (n = 13) and eYFP (n = 13) rats that had received prior CS-UCS conditioning without optogenetic stimulation. These rats were now allowed to lever press for the CS, during a session where each earned CS presentation was paired with laser stimulation.

**Figure 7.**
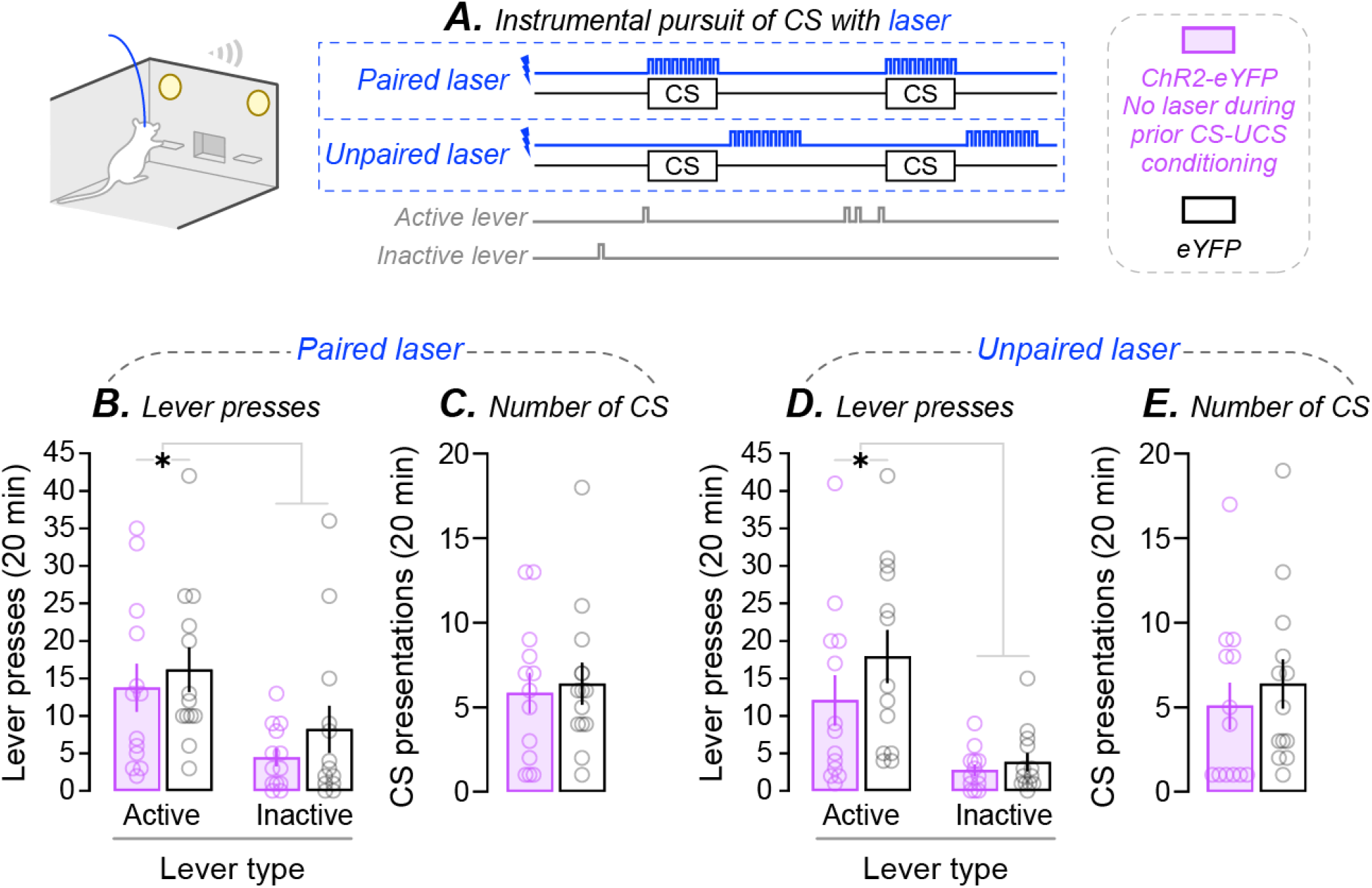
Optogenetic stimulation of BLA→NAc core neurons during instrumental conditioning does not alter the conditioned reinforcing properties of the CS. (**A**) We examined the effects of stimulating BLA→NAc core neurons during instrumental responding for CS using rats that had received initial CS-UCS conditioning without optogenetic stimulation (ChR2-eYFP, purple histograms; eYFP, white histograms). Rats could lever press on an active lever for CS presentation (RR2) and on an inactive lever. During the first instrumental conditioning session, laser stimulation was combined with CS presentation. During the second session, CS presentation and laser stimulation were unpaired (*i.e*., laser stimulation 3 seconds after CS presentation). When CS presentations were paired with laser stimulation, (**B**) both ChR2-eYFP and eYFP rats pressed more on the active than the inactive lever, with no group differences and (**C**) they obtained a comparable number of CS presentations during the session. Similarly, when laser stimulation was explicitly unpaired from CS presentation, (**D**) ChR2-eYFP and eYFP rats pressed more on the active than the inactive lever, with no group differences, and (**E**) they earned an equivalent number of CS presentations. ChR2-eYFP rats, n = 13; eYFP rats, n = 13. In (**B**, **D**); \**p* < 0.05, Lever type effect. Values in figures are means ± SEM. Individual data are shown on histograms.

#### Optogenetic stimulation of BLA→NAc core neurons during lever pressing for the CS does not influence instrumental responding

Fig. 7B shows that across groups, rats pressed more on the active versus inactive lever (Lever type effect, *F*_(1,24)_ = 17.16, *p* = 0.0004), and there were no group differences in this effect (*p* > 0.05). Thus, both groups performed an operant response reinforced solely by the CS, indicating that the CS has conditioned reinforcing effects, and these effects were similar across groups.

This is further highlighted by the observation that ChR2-eYFP and eYFP rats also earned a similar number of CS presentations (Fig. 7C; *p* > 0.05). To control for any potential non-specific effects of laser stimulation on behavior, we gave rats a final conditioned reinforcement test session during which we explicitly unpaired CS presentations and optogenetic stimulation of BLA→NAc core neurons, by applying this stimulation 3 seconds after each CS presentation. During this test, ChR2-eYFP and eYFP rats pressed more on the active versus inactive lever, without group differences (Fig. 7D; Lever type effect, *F*_(1,24)_ = 20.9, *p* = 0.0001). Both groups also earned a comparable number of CS presentations during the session (Fig. 7E; *p* > 0.05). These results indicate that optogenetic stimulation of BLA→NAc core neurons did not disrupt instrumental behavior.

Thus, optogenetic stimulation of BLA→NAc core neurons during instrumental responding for a CS does not influence the conditioned reinforcing properties of that CS. Taken together, the conditioned reinforcement tests (Figs. 6–7) show that activation of the BLA→NAc core pathway influences neither the attribution of conditioned reinforcing value to a CS nor the behavioral expression of conditioned reinforcement.

### Effects of optogenetic stimulation of BLA→NAc core neurons on extinction responding

Here we first determined whether optogenetic stimulation of BLA→NAc core neurons during initial CS-UCS conditioning influenced later conditioned approach behaviors triggered by CS presentations alone, without water (Fig. 8A). To this end, we used the following 3 groups of rats from Fig. 5: ChR2-eYFP rats that had received CS-UCS conditioning with (n = 11), or without (n = 6) CS-paired optogenetic stimulation, and eYFP rats (n = 8). All rats first received 2 reminder CS-UCS conditioning sessions, without laser stimulations (Fig. 8A; Sessions 1-2). They then received 2 extinction sessions during which the CS was presented alone (no water, and no laser stimulation; Sessions 3-4). ChR2-eYFP/no laser and eYFP rats showed similar responding during extinction tests and they were pooled (‘Controls’, n = 14). Next, we determined whether optogenetic stimulation of BLA→NAc core neurons *during* extinction influences Pavlovian conditioned approach behavior. To this end, the rats received 2 more reminder CS-UCS conditioning sessions (no laser stimulation; Sessions 5-6), followed by 2 final extinction sessions during which CS presentation (no water) was paired with optogenetic stimulation of BLA→NAc core neurons (Sessions 7-8).

**Figure 8.**
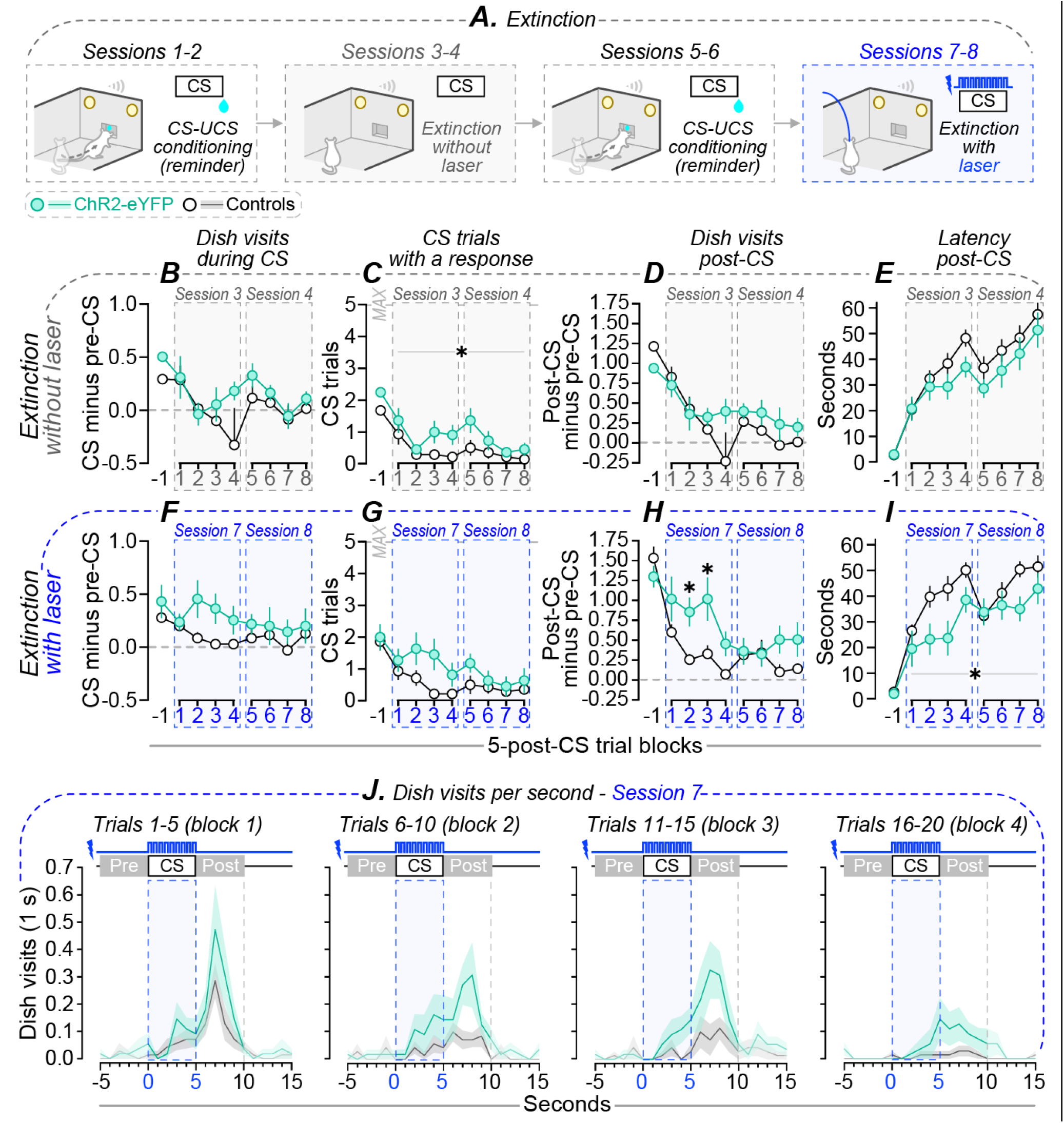
Optogenetic stimulation of BLA→NAc core neurons during extinction promotes CS-triggered increases in conditioned approach behavior. (**A**) We used the rats that had received initial CS-UCS conditioning with BLA→NAc core neuron stimulation to first determine whether this stimulation subsequently alters extinction learning. To do so, rats received 2 reminder CS-UCS conditioning sessions (Sessions 1-2; no laser stimulation), followed by 2 extinction sessions where the UCS was withheld (Sessions 3-4; no laser stimulation). Finally, we examined the effects of stimulating BLA→NAc core neurons *during* extinction learning. Thus, rats received 2 reminder CS-UCS conditioning sessions (Sessions 5-6; no laser stimulation), followed by 2 extinction sessions where each CS presentation was combined with laser stimulation (Sessions 7-8). We examined responses across 5-trial blocks during the extinction sessions. Trial block ‘−1’ corresponds to the mean response per 5-trial block on the conditioning session prior to extinction, when the CS was still followed by the water UCS. During the extinction tests without laser stimulation (Sessions 3-4), rats reduced their number of (**B**) dish visits during CS presentation (number of dish visits during the 5-s CS minus the number of dish visits 5 s prior to each CS, pre-CS), (**C**) CS trials with at least one dish visit during the CS, (**D**) dish visits following CS presentation (number of dish visits during the 5-s period after the CS, post-CS, minus the number of dish visits during the pre-CS), and (**E**) they increased their latency to enter the water dish after CS presentation. ChR2-eYFP and eYFP rats showed a similar decrease in CS-evoked responses, except that (**C**) ChR2-eYFP rats responded to a greater number of CS trials compared to controls. During the extinction sessions where each CS presentation was now combined with laser stimulation (Sessions 7-8), rats across groups (**F**) maintained a stable number of dish visits during the CS across trials, however they (**G**) decreased the number of CS trials they responded to with at least one dish visit, (**H**) decreased the number of post-CS dish visits and (**I**) increased their post-CS latency. ChR2-eYFP rats made (**H**) more post-CS dish visits and (**I**) showed a reduced latency to enter the water dish after CS presentation compared to controls. (**J**) Water dish visits per second before, during and after CS presentations during Session 7, with averaged responses of trials 1-5 (block 1), 6-10 (block 2), 11-15 (block 3) and 16-20 (block 4). ChR2-eYFP rats, n = 11; controls, n = 14. \**p* < 0.05; in (**C**, **I**), Group effect; in (**H**), relative to controls during the same 5-trial block. Values in figures are means ± SEM.

#### Effects of optogenetic stimulation of BLA→NAc core neurons during initial Pavlovian conditioning on subsequent extinction behavior

Each extinction session contained 20 CS trials. We analysed the effects of optogenetic stimulation on CS-triggered conditioned approach behavior in 5-trial blocks across each session. In Figs. 8B-I, trial block ‘−1’ represents the mean response per 5-trial block on the Pavlovian conditioning session prior to extinction, when each CS was still followed by water. During extinction sessions 3-4, which occurred without laser stimulation, ChR2-eYFP rats and controls reduced the number of dish visits they made during the CS, the number of CS trials they responded to with at least one water dish visit, the number of post-CS dish visits, and the latency to enter the water dish after CS presentation (Trial effect; Fig. 8B, *F*_(7,161)_ = 2.19, *p* = 0.038; Fig. 8C, *F*_(7,161)_ = 21.62, *p* < 0.0001; Fig. 8D, *F*_(7,161)_ = 4.46, *p* = 0.0001; Fig. 8E, *F*_(7,161)_ = 10.93, *p* < 0.0001). Thus, when water no longer followed each CS presentation, rats in both groups reduced the frequency and speed of visits to the water dish.

ChR2-eYFP rats responded to more CS trials compared to controls (Fig. 8C; Group effect, *F*_(1,23)_ = 4.6, *p* = 0.04). However, there were no group differences in the number of dish visits during or after the CS, or in the post-CS latency (Figs. 8B and 8D-E; all *P*’s > 0.05). Thus, stimulating BLA→NAc core neurons during initial CS-UCS conditioning had no effect on most measures of CS-evoked conditioned approach behavior during later extinction sessions.

#### Optogenetic stimulation of BLA→NAc core neurons during extinction sessions reduces extinction responding

During extinction sessions 7-8, each CS presentation was now paired with laser stimulation. The number of dish visits made during CS presentations remained stable across trials (Fig. 8F; no Trial effect, no Group effect; all *P*’s > 0.05). However, across groups, the rats significantly reduced both the number of CS trials they responded to and their post-CS water dish visits, and they also increased their post-CS latency (Trial effect; Fig. 8G, *F*_(7,161)_ = 3.79, *p* = 0.0008; Fig. 8H, *F*_(7,161)_ = 5.91, *p* < 0.0001; Fig. 8I, *F*_(7,161)_ = 7.84, *p* < 0.0001; no Group effects, all *P*’s > 0.05). Thus, when the CS was presented alone, the rats reduced their conditioned approach responses. There were also group differences in responding. First, ChR2-eYFP rats made significantly more post-CS dish visits than control rats did (Fig. 8H; Group × Trial interaction, *F*_(7,161)_ = 2.11, *p* = 0.046; Group effect, *F*_(1,23)_ = 6.5, *p* = 0.02), in particular during Session 7 (ChR2-eYFP rats > controls; trial block 2, *p* = 0.04; trial block 3, *p* = 0.01). Second, ChR2-eYFP rats visited the water dish sooner after CS presentation compared to controls (Fig. 8I; Group effect, *F*_(1,23)_ = 6.32, *p* = 0.02). Thus, when the CS is presented without the water UCS, pairing CS presentations with optogenetic stimulation of BLA→NAc core neurons increases CS-triggered conditioned approach behaviors. This effect is further highlighted in Fig. 8J, which illustrates dish visits per second across the 20 CS trials of extinction session 7, during which each CS was paired with laser stimulation. ChR2-eYFP and control rats showed similar responding *during* CS presentations. However, immediately after CS presentation, ChR2-eYFP rats approached the water dish at a greater rate than control rats did, despite the absence of the water UCS.

In summary, optogenetic stimulation of BLA→NAc core neurons during prior Pavlovian CS-UCS conditioning does not significantly influence later extinction responding, save for an effect on the number of CS trials with at least one dish visit. However, stimulating these neurons during extinction sessions increases both the frequency and speed of CS-evoked conditioned approach responses, even though the CS was presented alone, without the water UCS.

## DISCUSSION

The BLA modulates both conditioned approach behavior triggered by an appetitive CS and the instrumental pursuit of such CS (Cador et al., 1989; Burns et al., 1994; Servonnet et al., 2020). Here we evaluated the contributions of BLA→NAc core neurons to these effects. During Pavlovian CS-water conditioning, optogenetic activation of BLA→NAc core neurons enhanced CS-triggered conditioned approach behavior to the site of water delivery. Stimulating BLA→NAc core neurons also enhanced CS-evoked conditioned approach responses under extinction conditions, when the water UCS no longer followed the CS. In parallel, rats did not self-administer BLA→NAc core neuron stimulations, indicating that these neurons do not carry a primary reward signal. Finally, activation of BLA→NAc core neurons either during initial Pavlovian CS-UCS conditioning or during subsequent lever pressing for the CS did not alter instrumental responding for the CS. This suggests that activity in BLA→NAc core neurons does not influence the conditioned reinforcing value of an appetitive CS. Together, the results suggest that activation of BLA→NAc core neurons facilitate cue-induced control over behavior by promoting conditioned expectation of reward, without affecting the operant pursuit of reward cues.

### Stimulation of BLA→NAc core neurons promotes CS-triggered anticipation of reward

During CS-UCS conditioning, rats that received CS-paired optogenetic stimulation of BLA→NAc core neurons more readily developed conditioned approach behavior. Compared to controls, these rats showed greater CS-triggered visits to the water dish, and they also developed this conditioned response earlier across sessions. Previous work shows that photoinhibition of BLA→NAc projections (without distinguishing between the core and shell subregions) supresses Pavlovian conditioned approach behaviors (Stuber et al., 2011), suggesting that activity in BLA→NAc neurons is necessary for this CS effect. Our findings extend this observation by showing that activity in BLA→NAc core neurons is sufficient to potentiate CS-evoked Pavlovian approach. Conversely, optogenetic stimulation of BLA→NAc shell neurons *reduces* CS-triggered conditioned approach behavior (Millan et al., 2017). This suggests that BLA projections to different NAc subdivisions play distinct roles in CS-UCS conditioning, with projections to the core promoting conditioned approach behavior, and projections to the shell suppressing this behavior. Thus, when animals encounter reward-predictive cues, excitatory signals from the BLA to the NAc core enhance appetitive approach responses, suggesting increased conditioned anticipation of reward (Tolman, 1932; Hearst and Jenkins, 1974; Cardinal et al., 2002).

Optogenetic stimulation of BLA→NAc core neurons also influenced conditioned approach responses under extinction. Rats that had received CS-paired optogenetic stimulation of BLA→NAc core neurons during initial Pavlovian conditioning later showed normal extinction responding—*i.e*., they decreased their CS-triggered conditioned approach responses in the absence of water reward. However, stimulating BLA→NAc core neurons during extinction increased CS-evoked conditioned responses, suggesting persistent conditioned anticipation of the water reward, even when this reward no longer followed the CS. Prior work shows that activity in BLA→NAc core neurons is required for CS-induced reinstatement of instrumental responding for reward (Stefanik and Kalivas, 2013). Together, this and our finding suggest that under extinction conditions, BLA→NAc core neurons regulate CS-triggered increases in both instrumental responding for reward and Pavlovian conditioned approach responses.

Activating BLA→NAc core neurons might promote CS-evoked conditioned approach responses via several mechanisms. First, activation of BLA→NAc core neurons might increase the appetitive value of the water UCS *per se*, without necessarily influencing CS properties. However, this seems unlikely. In a study where mice were trained to associate a CS with sucrose reward, photoinhibition of BLA→NAc core neurons *after* CS presentation—when mice would be consuming the sucrose—did not influence conditioned approach behavior (Namburi et al., 2015). This suggests that in a conditioning paradigm, these neurons do not mediate the appetitive properties of the primary reward. Second, the water dish was also a CS in our experiments, and activating BLA→NAc core neurons could have increased water-dish entries by potentiating the dish’s incentive value, independent of any effects on the discrete light-tone CS. However, this is unlikely because BLA→NAc core neuron stimulation increased dish visits specifically during CS-UCS trials, without influencing dish entries at other times during conditioning sessions. Third, stimulation might enhance CS-evoked representation of specific characteristics of the associated reward. This can include the sensory properties of the associated water reward and/or its appetitive value (Cardinal et al., 2002), thus potentiating CS-triggered increases in the incentive urge to consume the water (Weingarten, 1983). Finally, stimulation of BLA→NAc core neurons might also enhance the emotional tone that is tagged to the CS, thereby enhancing CS-evoked expectation of the associated reward (Tolman, 1932; Hearst and Jenkins, 1974; Cardinal et al., 2002).

### Stimulation of BLA→NAc core neurons is not intrinsically reinforcing and does not influence the instrumental pursuit of a CS

Similar to our previous finding that rats do not self-administer optogenetic stimulations of BLA neurons (Servonnet et al., 2020), here we found that rats do not reliably self-administer optogenetic stimulations of BLA→NAc core neurons. This is in apparent contrast with previous studies showing that mice self-stimulate optogenetic stimulation of BLA→NAc neurons (Stuber et al., 2011; Namburi et al., 2015). This discrepancy could involve the use of rats versus mice, but also the NAc subregion targeted. Stuber et al. (2011) and Namburi et al. (2015) did not distinguish between BLA inputs to NAc core versus shell, while here we targeted BLA→NAc core inputs specifically. Interestingly, mice self-administer optogenetic stimulation of BLA→NAc *shell* neurons (Britt et al., 2012). Together with our findings, this suggests that BLA projections to the NAc shell but not core carry a primary reward signal supporting self-stimulation.

Stimulating BLA→NAc core neurons did not influence the conditioned reinforcing properties of a reward-predictive CS. Rats were allowed to acquire a new lever-pressing response reinforced solely by the CS (without water). Both controls and rats that had received CS-paired optogenetic stimulations either during previous Pavlovian conditioning or *during* lever-pressing learned this new instrumental response, indicating that the CS had acquired conditioned reinforcing properties. However, there were no group differences in lever responding for the CS, suggesting that stimulation of BLA→NAc neurons changes neither the acquisition nor the expression of CS reinforcing value. This extends previous findings showing that the BLA is not required for the ability of intra-NAc core amphetamine to potentiate lever responding for an appetitive CS (Cador et al., 1989; Burns et al., 1993), suggesting that BLA-NAc core interaction is not necessary for this cue-controlled behavior.

Recently, we found that optogenetic stimulation of BLA neurons without targeting specific projections potentiates the instrumental pursuit of a reward-predictive CS (Servonnet et al., 2020). Together with our present results, this suggests that the BLA mediates the conditioned reinforcing properties of CS through recruitment of a neural pathway that does not involve direct projections to the NAc core. Future studies can examine the potential contributions of BLA projections to the NAc shell or prefrontal cortex to this effect (Mashhoon et al., 2010; Stefanik and Kalivas, 2013; Keistler et al., 2017; Lichtenberg et al., 2017; Lichtenberg et al., 2021). While the present findings show that activation of BLA→NAc core neurons does not influence the conditioned reinforcing properties of discrete CS, these neurons might still regulate these properties of reward-associated *contexts*. For instance, the expression of conditioned preference for a cocaine-paired context increases Fos expression in BLA→NAc core neurons, suggesting increased recruitment of these neurons [(Miller and Marshall, 2005) see also (Everitt et al., 1991)].

### Considerations

We only used male rats, and there are limitations to generalizing results obtained with males to females (Prendergast et al., 2014; Becker and Koob, 2016; Shansky, 2019; Shansky and Murphy, 2021). Our findings lay some of the initial groundwork for similar investigations across the sexes. In addition, optogenetic stimulation of neuronal terminals can evoke antidromic action potentials (Fig. 3A). Thus, the observed effects could have been influenced by activation of BLA→NAc core neuron collaterals to other brain regions, such as the NAc shell, the prelimbic cortex and/or the anterior insula (Shinonaga et al., 1994; Beyeler et al., 2016; Puaud et al., 2021). This can be addressed by extending the present work with photoinhibition methods. Importantly, this issue does not change our main conclusion that stimulation of BLA terminals in the NAc core has dissociable effects on CS-triggered anticipation of reward versus conditioned reinforcement.

### Conclusions

Our findings show that pairing CS presentation with activation of BLA→NAc core neurons enhances CS-triggered increases in approach to the site of reward delivery. This enhancement is observed both during initial CS-UCS conditioning and under extinction conditions, when rats encounter the CS without the primary reward. This suggests that increased activity of BLA→NAc core neurons promotes a cue-induced anticipatory state, guiding appetitive behavior in response to environmental change. In parallel, activating BLA→NAc core neurons does not influence the instrumental pursuit of a reward-paired CS, indicating no effect on the motivational state of ‘wanting’ for the reward cues (Robinson and Berridge, 1993). Extending previous findings (Ambroggi et al., 2008; Stefanik and Kalivas, 2013; Puaud et al., 2021), we conclude that while increased activity in BLA→NAc core neurons does not change the ability of CS to reinforce the learning of new reward-seeking actions, it enhances CS-driven reward anticipation, thereby preparing animals to engage with forthcoming rewards.

## ACKNOWLEDGEMENTS

We thank Dr. Giovanni Hernandez and Charles Ducrot for valuable advice on electrophysiological recordings and immunohistochemistry, respectively. We thank Dr Jonathan Britt for technical advice on *in vivo* optogenetics. We thank Dr Mike JF Robinson for comments on an earlier manuscript draft. This work was supported by grants from the National Science and Engineering Research Council of Canada (grant number 355923) and the Canada Foundation for Innovation to ANS (grant number 24326). ANS holds a salary award from the Fonds de la Recherche du Québec-Santé (grant number 28988).

## Notes

**CONFLICT OF INTEREST** The authors declare no competing financial interests.

### Competing Interest Statement

The authors have declared no competing interest.

